# Pancreatic mu opioid receptors regulate metabolism and ingestive behaviors

**DOI:** 10.64898/2026.05.21.726374

**Authors:** Diego A. De Gregorio, Daniel C. Castro

## Abstract

The mu opioid receptor system has long been recognized for its role in regulating ingestive behaviors and food palatability. However, beyond their robust expression in the brain, opioid receptors are also found throughout the body, including the stomach, intestines, and pancreas. Previous studies suggest that opioids may directly regulate glycemia through actions in the endocrine pancreas, but whether these metabolic effects are linked to their influence on ingestive behavior remains unclear. Here, we used metabolic, ingestive, and genetic approaches to determine how the mu opioid receptor (MOPR) regulates in vivo phenotypes via its actions in endocrine pancreas. Glucose and insulin tolerance tests in male *Oprm1* KO mice had enhanced glucose tolerance and insulin sensitivity. Conditional deletion of MOPRs in alpha cells had no effect. To test if loss of MOPRs influence overt ingestive behaviors, we measured 24-hour ad libitum food consumption using Feeding Experimental Devices (FEDs) in wild type, *Oprm1* knockout (MOPR-deficient) and *GCG*-Cre x *Oprm1*^*fl/fl*^ mice (10-12 weeks old). We found that *Oprm1* KO male, but not female, mice exhibited increased mass and food intake, and dysregulated ingestive microstructures relative to wild type controls. Conditional deletion of MOPRs in alpha cells only recapitulated some of the phenotypes related to ingestive microstructure. These findings suggest that MOPR regulation of ingestive behaviors extends beyond typical neural circuits of reward to include peripheral metabolic organ mechanisms.

**Highlights:**

- Male *Oprm1* knockout mice show increased body mass and food intake
- Male, but not female, *Oprm1* knockout mice have enhanced glucose tolerance and insulin sensitivity
- MOPR deletion in pancreatic alpha cells did not alter glucose metabolism
- MOPR deletion in alpha cells increased meal frequency but reduced meal size in males
- MOPR deletion in alpha cells only reduced meal size in females

## Introduction

Obesity, diabetes, and metabolic disorders are among the most pressing and taxing public health challenges in the United States. Over 42% of U.S. adults are classified as obese, and nearly 12% live with diabetes^1^. These conditions significantly increase the risk of comorbidities such as cardiovascular disease^2,3^ and peripheral neuropathy^4–6^, the latter of which is often treated with exogenous opioid drugs like tramadol^7,8^, oxycodone^9,10^, and fentanyl.^11,12^ These drugs act primarily through the μ-opioid receptor (MOPR), a G-protein-coupled receptor involved in both pain modulation^13–17^ and central reward processing.^18–2021–25,20^

MOPRs play a well-established role in modulating ingestive behaviors and the hedonic aspects of food consumption^26,19^. Activation of MOPRs in brain regions such as the nucleus accumbens enhances the palatability of high-fat and high-sugar foods and increases food intake^20,27,28,28,29^, while regions like the hypothalamus and insular cortex integrate opioid-mediated signals related to hunger, satiety, and reward^30,31,19^. Importantly, these central effects may influence not only how much is eaten but also how meals are structured, including meal size, frequency, and ingestive circadian timing factors.^15,32^

In addition to its central roles, MOPR is also expressed peripherally, including in the stomach^33,34^, intestines^35,36^, and endocrine pancreas^37,38^. Within the endocrine pancreas, MOPRs are present across all key islet cell populations, including α-cells, β-cells and other hormone-secreting subtype cells^39^. Paradoxically, constitutive deletion of MOPRs results in greater adiposity, but enhances glucose tolerance and insulin secretion. These data suggest that the role of MOPRs on regulating metabolism may be more nuanced that previously appreciated. Support for this conclusion was recently published wherein the glucagon producing α-cells were identified as a particularly important site for MOPR action. Specifically, disruption of MOPRs via pharmacological antagonism in human islets, or selective genetic deletion from α-cells in mouse islets, increased glucagon secretion. Additionally, while α-cell MOPR knockout mice had normal baseline glucose and insulin tolerance, exposure to a mild high fat diet revealed a higher propensity for developing a Type 2 diabetes-like phenotype. Similarly, patients with Type 2 diabetes and obese mice were found to have downregulated MOPR expression, indicating a dynamic relationship between MOPRs, pancreatic islets, and metabolic phenotypes. Altogether, while these results highlight an important role for MOPRs on alpha cells, it remains unknown whether or how modulation of MOPRs on α-cells ultimately impacts overt ingestive behaviors.^38^

Therefore, to characterize the role of MOPRs in metabolic and ingestive regulation, we compared the effects of global MOPR deletion and selective deletion of MOPRs from pancreatic α-cells. Using in vivo assessments of glucose homeostasis, alongside analyses of ingestive microstructure in male and female mice, we found that global deletion of MOPRs enhanced glucose and insulin tolerance and increased meal frequency, particularly during the dark cycle, in male but not female mice. In contrast, selective deletion of MOPRs from α-cells did not reproduce any metabolic phenotype observed in global knockouts but did alter ingestive microstructure by increasing meal frequency and reducing meal size, primarily in males. Together, these findings suggest that MOPRs regulate metabolism and feeding behavior through both central and peripheral mechanisms, with α-cell MOPRs contributing specifically to meal patterning rather than global metabolic control under otherwise healthy conditions.

## 2. Material and methods

### 2.1 Animals

Adult (10-14 weeks) male and female (18-35g) wild type, *Oprm1* KO, and *Oprm1*^fl/fl^ x *GCG*-Cre mice were group housed, given access to food pellets and water ad libitum, and maintained on a 12□h:12□h light:dark cycle (lights on at 06:00). All mice were kept in a sound-attenuated, isolated holding facility one week prior and throughout the duration of the behavioral assays to minimize stress. *Oprm1* KO mice were compared with age matched wild type mice. *Oprm1*^fl/fl^ x *GCG*-Cre+ mice were compared with age matched littermate Cre-mice. Unless otherwise noted, mice had ad libitum access to food and water. Any variation from these approaches was owing to behavioral attrition. The mice were bred at Washington University in Saint Louis. All mice were drug and test naive, individually assigned to specific experiments as described, and not involved with other experimental procedures. All mice were monitored for health status daily and before experimentation for the entirety of the study. All procedures were approved by the Animal Care and Use Committee of Washington University in Saint Louis, Animal Care and Use Committee of Washington University in Saint Louis and conformed to US National Institutes of Health guidelines.

### 2.4 Glucose and Insulin Tolerance Tests

Adult male and female C57BL/6J, *Oprm1* KO, and *Oprm1*^*fl/fl*^ *x GCG-Cre* mice (8–12 weeks old) were habituated to the experimenter the day prior to each test. Mice were fasted for 6 to 8 hours before glucose tolerance testing (GTT) and insulin tolerance testing (ITT). GTT and ITT were counterbalanced across mice and separated by at least one week. For GTT, mice received an intraperitoneal injection of 20% D-glucose (1g/kg body weight). For ITT, mice received an intraperitoneal injection of human regular insulin (0.75 U/kg; Lilly, Indianapolis). Blood samples (5 μL) were collected via tail nick at 0, 15-, 30-, 60-, and 120-minutes post-injection, and glucose concentrations were measured using a handheld glucometer (Arkray USA 761100, Glucocard® Vital™). All tests were performed during the light cycle between 9:00 AM and 6:00 PM.

### 2.5 Biochemical Assays of Blood Serum

Blood was collected from the tail vein after a 4-6-hour fast using a heparinized capillary tube. Samples were transferred to 1.5mL microcentrifuge tubes submerged in an ice bucket immediately after blood was taken. We then centrifuged at 6000rpm for 15 minutes at 4°C to separate plasma. The supernatant was carefully collected and stored at −80°C until analysis. Concentrations of triglycerides, total cholesterol, glucose, and free fatty acids (FFAs) were quantified using colorimetric assay kits according to the manufacturer’s instructions. All assays were performed by Animal Models Phenotyping Core at Washington University in Saint Louis and absorbance was read using a microplate spectrophotometer (e.g., SpectraMax iD3, Molecular Devices).

### 2.6 Food intake test

Mice were kept singly housed for four consecutive days with given access to food pellets and water ad libitum and maintained on a 12□h:12□h light:dark cycle (lights on at 06:00). All mice were kept in a sound-attenuated, isolated holding facility one week prior and throughout the duration of the behavioral assays to minimize stress. Mice were weighed in each test date. After the 24-hour habituation period, unobtrusive monitoring of food intake behavior (Dustless Precision Pellets® Rodent, Grain-Based, BioServ) was measured for three consecutive days using Feeding Experimental Devices (FEDs). Using an Arduino, each FED’s magazine and nose ports were outfitted with IR beams to capture subsecond behavioral engagement. Total pellets consumed, number of meals, and meal size were quantified. A meal was operationally defined as any bout of one pellet or more with a maximum interpellet interval of one minute. Data was collected in the form of an excel sheet and data was transferred via an integrated SD card using the fed3viz software. For data analysis, a MATLAB script was used, and data was plotted using Prism. All mice were drug and test naive, individually assigned to specific experiments as described, and not involved with other experimental procedures. All mice were monitored for health status daily and before experimentation for the entirety of the study. All procedures were approved by the Animal Care and Use Committee of Washington University in Saint Louis, Animal Care and Use Committee of Washington University in Saint Louis and conformed to US National Institutes of Health guidelines.

## 3. Results

Male and female mice were analyzed separately to assess potential sex differences across behavioral or metabolic phenotypes.

### 3.1.1 MOPR knockout metabolic phenotypes

Male *Oprm1* KO mice had significantly enhanced glucose tolerance relative to wild type male mice in an IP glucose tolerance test (Time x Genotype: F (2.171, 39.07) = 9.662, p = 0.0003). Baseline blood glucose did not differ between genotype, but enhancement was observed all timepoints tested after glucose administration (wild type vs KO 0 min t(17.60) = 1.562, p = 0.1361; wild type vs KO 15 min t(16.18) = 5.073, p = 0.0001; wild type vs KO 30 min t(17.33) = 5.587, p < 0.0001; wild type vs KO 60 min t(17.65) = 2.709, p = 0.0145; wild type vs KO 90 min t(17.42) = 2.884, p = 0.0101; wild type vs KO 120 min t(15.91) = 3.040, p = 0.0078). A similar phenotype was observed with insulin tolerance, such that male KO mice had increased insulin sensitivity during an insulin tolerance test, albeit baseline measurements were again not different (Genotype: F (1, 17) = 23.80, p = 0.0001; wild type vs KO 0 min t(16.63) = 1.981, p = 0.0644; wild type vs KO 15 min t(9.085) = 2.606, p = 0.0282; wild type vs KO 30 min t(11.42) = 4.509, p = 0.0008; wild type vs KO 60 min t(14.59) = 5.126, p = 0.0001; wild type vs KO 90 min t(12.11) = 3.114, p = 0.0089). Finally, we tested whether there were any differences between various blood serum metabolic biomarkers, including cholesterol, free fatty acids, or triglycerides. We did not observe any baseline differences between male wild type or KO mice (Biomarker x Genotype: F (1.128, 19.18) = 1.952, p = 0.1783; Cholesterol: t(16.97) = 0.2940, p = 0.9973; Triglycerides: t(16.88) = 0.8410, p = 0.8805; FFA: t(16.82) = 0.7501, p = 0.9172).

Unlike male KO mice, glucose tolerance testing in female *Oprm1* KO mice revealed no phenotype relative to wild type females (Genotype: F (1, 18) = 0.7480, p = P=0.3985). Similarly, while all female mice showed robust insulin sensitivity during insulin tolerance testing, this effect was not different between wild type and KO mice (Genotype: F (1, 18) = 3.335, p = 0.0845). Lastly, we did not observe any differences in blood serum biomarkers (Biomarker x Genotype: F (1.257, 22.62) = 1.149, p = P=0.3099; Cholesterol: t(17.85) = 0.2964, p = 0.9972; Triglycerides: t(16.55) = 0.6465, p = 0.9499; FFA: t(17.78) = 1.579, p = 0.4324). Together, these results indicate that constitutive MOPR knockout has a greater impact on male glycemic responses compared to female responses, and that male mice have both increased responsivity (glucose tolerance) and sensitivity (insulin tolerance). However, other metabolic biomarkers were unaffected, suggesting that MOPRs ultimate effects on metabolism may be driven by glycemic regulators.

### 3.1.2. MOPR knockout ingestive phenotypes

Male *Oprm1* KO mice weighed significantly more relative to wild type males (median KO = 29, median wild type = 25; U = 9, p < 0.0001). Correspondingly, KO males also ate more pellets over the course of the three day test (median KO = 718, median wild type = 621; U = 22.5, p < 0.0001), however there was no correlation between weight and intake among KO male mice, indicating caloric demand was insufficient to explain the phenotype (r = 0.1163, n = 15, p = 0.6798). Further analyses revealed that while both KO and wild type males increased consumption during the dark cycle versus the light cycle (F (5, 140) = 9.879; p < 0.0001; wild type ZT2 vs ZT14 t(14) = 2.843, p = 0.0026; KO ZT2 vs ZT14 t(14) = 12.05, p < 0.0001), this effect was amplified in KO males (wild type vs KO ZT10 t(27.94) = 4.144, p = 0.0003; ZT14 t(23.16) = 5.064, p < 0.0001; ZT18 t(27.06) = 3.106, p = 0.0044). Next, we sought to determine whether wild type and KO male mice differed in ingestive microstructure, specifically evaluating the number of meals during the testing period, as well as meal size. Overall, there was no difference in meal number between wild type and KO male mice (median wild type = 108, median KO = 138; U = 67, p = 0.0597). However, meal number did change across the light/dark cycle, with both wild type and KO male mice increasing meal number during the dark cycle (F (5, 140) = 3.313. p = 0.0074; wild type ZT6 vs ZT18 t(14) = 2.575, p = 0.0268; KO ZT6 vs ZT18 t(14) =8.676, p < 0.0001). Follow up analyses revealed that when time was taken into account, KO mice ate significantly more meals than wild type males, but only during the dark phase (wild type vs KO ZT18 t(27.88) = 2.572, p = 0.0157). In contrast, meal size was stable throughout the testing period, and did not differ between wild type and knockout mice (median KO = 3.959, median wild type = 4.734; U = 95, p = 0.4864).

Unlike male mice, *Oprm1* KO female mice did not weigh more than wild type female mice (median KO = 22, median wild type = 21; U = 73, p = 0.0865). KO females also consumed a similar number of pellets as wild type females (median KO = 663, median wild type = 621; U = 94, p = 0.461), and while both KO and wild type female mice increased consumption during the dark phase relative to the light phase (F (3.933, 110.1) = 45.94, p < 0.0001; wild type ZT2 vs ZT14 t(14) = 8.844, p < 0.0001; KO ZT2 vs ZT14 t(14) = 8.652, p < 0.0001), there was no difference between genotypes (F (5, 140) = 1.566, p = 0.1735). Despite no overt differences between KO and wild type females, KO females did show some minor phenotypes related to ingestive microstructure. Specifically, KO females ate more meals compared to wild type females, but only at the transition from the light phase to the dark phase (median KO = 114, median wild type = 106; U = 94, p = 0.2452; F (5, 140) = 4.039, p = 0.0019; wild type vs KO ZT10 t(27.91) = 2.828, p = 0.0099; wild type vs KO ZT14 t(25.55) = 2.017, p = 0.0544). Like males, wild type and KO female mice did not differ in meal size (median KO = 3.186, median wild type = 4.479; U = 76, p = 0.1370). Altogether, while female KO mice had fewer and less robust ingestive phenotype relative to wild type females, there are a few notable similarities between KO females and males.

### 3.2.1 Conditional alpha cell MOPR knockout metabolic phenotypes

Male *Oprm1*^*fl/fl*^ x *GCG-*Cre*-* and *Oprm1*^*fl/fl*^ x *GCG-*Cre*+* mice had similar glucose tolerance values at all timepoints tested (Genotype: F (1, 18) = 0.1047, p = 0.75). Blood glucose values followed typical patterns with a peak at 30 minutes post-injection (Cre-0 vs 30 min, t(9) = 7.779, p = 0.0004; Cre+ 0 vs 30 min, t(9) = 11.76, p < 0.0001), and returning to baseline by 120 minutes injection (Cre-0 vs 120 min, t(9) = 0.5066, p > 0.99; Cre+ 0 vs 120 min, t(9) = 0.7640, p > 0.99). Like glucose tolerance, male Cre- and Cre+ mice had nearly identical insulin tolerance values (Genotype: F (1, 18) = 0.03584, p = 0.8520) and followed normal blood glucose trajectories in response to insulin (Time: F (2.600, 46.81) = 43.08, p < 0.0001). Finally, male Cre- and Cre+ mice did not show any differences between the blood serum biomarkers tested, including cholesterol, free fatty acids, or triglycerides (Biomarker x Genotype: F (3, 57) = 3.038, p = 0.0363; Cholesterol: t(76.00) = 0.2186, p = 0.9991; Triglycerides: t(76.00) = 1.491, p = 0.4530; FFA: t(76.00) = 0.01559, p > 0.9999).

Like male mice, female Cre- and Cre+ mice did not differ in glucose tolerance tests (Genotype: F (1, 18) = 0.1047, p = 0.75), showing blood glucose trajectories (Time: F (1.841, 33.13) = 69.61, p < 0.0001). A similar phenotype was observed with insulin tolerance testing (Genotype: F (1, 13) = 0.02083, p = 0.8875; Time: F (2.008, 26.11) = 37.93, p < 0.0001). Lastly, no differences in blood serum biomarkers were observed between female Cre- and Cre+ mice (Biomarker x Genotype: F (3, 51) = 0.06739, p = 0.9770; Cholesterol: t(68.00) = 0.3421, p = 0.9949; Triglycerides: t(68.00) = 0.1663, p = 0.9997; FFA: t(68.00) = 0.02200, p > 0.9999).

### 3.3.2 Conditional alpha cell MOPR knockout ingestive phenotypes

Male Cre+ mice did not differ in weight relative to Cre-males (median Cre+ = 27, median Cre- = 27; U = 95, p = 0.4648), nor did they differ in total amount of pellets consumed (median Cre+ = 613, median Cre- = 611; U = 106, p = 0.7984). Both Cre+ and Cre-male mice showed typical circadian ingestive behaviors, increasing their intake during the night cycle relative to the light cycle (F (3.438, 96.27) = 34.70; p < 0.0001; Cre-ZT2 vs ZT14 t(14) = 8.86, p = 0.0002; Cre+ ZT2 vs ZT14 t(14) = 12.29, p < 0.0001), but no interaction by genotype was detected at any time point (F (3.438, 96.27) = 1.178, p = 0.3243). Although no major differences were observed in total intake, Cre+ mice did display unique ingestive microstructure phenotypes. First, Cre+ male mice ate significantly more meals than Cre-males (median Cre+ = 144, median Cre- = 92; U = 57, p = 0.0209). This effect was most robust during the light phase, but a general trend can be observed across the light/dark cycle (F (2.399, 67.17) = 22.39, p < 0.0001; Cre-vs Cre+ ZT2 t(23.46) = 2.9, p = 0.008; Cre-vs Cre+ ZT10 t(27.99) = 3.244, p = 0.003). This increase in meals was offset by Cre+ male mice having smaller meals relative to Cre-males (median Cre+ = 4.022, median Cre- = 5.619; U = 50, p = 0.0086). Like the increased meals, this appeared to be a relatively stable trait across the light/dark cycle (by Genotype: F (1, 28) = 9.595, p = 0.0044; Cre-vs Cre+ ZT2 t(27.96) = 3.515, p = 0.0015; ZT6 t(25.85) = 1.939, p = 0.0635; ZT10 t(22.32) = 2.671, p = 0.0139; ZT14 t(25.94) = 2.619, p = 0.0145; ZT18 t(27.45) = 2.178, p = 0.0381; ZT22 t(27.33) = 2.802, p = 0.0092). In sum, these data indicate that while Cre+ mice share some ingestive phenotypic traits with constitutive KO males (e.g., increase meals), selective deletion of MOPRs from alpha cells appears to drive a distinct suite of ingestive microstructural phenotypes, highlighting a unique contribution of MOPRs to this cell type and ultimate ingestive behaviors.

Female Cre+ mice were largely indistinguishable from Cre-females. There were no differences in weight, overall intake, or number of meals between genotypes (Weight: median Cre+ = 20, median Cre- = 22; U = 67.5, p = 0.0615; intake: median Cre+ = 527.5, median Cre- = 524; U = 89, p = 0.4976; meal number: median Cre+ = 122.5, median Cre- = 101; U = 75.5, p = 0.2049). However, like Cre+ males, Cre+ female mice consistently had smaller meal sizes (median Cre+ = 3.142, median Cre- = 3.951, U = 44, p = 0.0068), regardless of the day/night cycle (by Genotype: F (1, 27) = 7.901, p = 0.0091; Time x Genotype: F (3.589, 96.89) = 2.119, p = 0.0913). In sum, male and female Cre+ mice showed decreased meal sizes, but this decrease was largely overcome by compensatory responses in other ingestive microstructures, resulting no detectible differences in weight or overall consumption.

## 4.1 Discussion

In this study we sought to determine how mu opioid receptors regulate metabolic and ingestive phenotypes, and to assess to what degree these phenotypes are driven by MOPRs on pancreatic α-cells. Using a global knockout mouse line, we first corroborate previous reports that loss of mu opioid receptors in male mice enhances glucose tolerance and increases insulin sensitivity^40^. This metabolic phenotype appeared to be exclusive to glycemia, as other metabolic biomarkers did not differ between knockout and wild type male mice. Efficient glucose handling can influence feeding by stabilizing energy availability^41^, potentially altering the drive to initiate meals^42,43^. To assess this, we continuously monitored chow intake for three days using home cage FED devices. We found that male knockout mice weighed significantly more than their wild type counterparts and correspondingly ate more pellets over the three test days. While overall ingestive microstructure did not differ between knockout and wild type mice, further analyses revealed that knockout mice engaged in more frequent meals during the early phase of the dark cycle. Meal size, by contrast, was consistent across the light/dark cycle, and was not different from wild type mice. This could suggest that the enhanced metabolic phenotypes observed in male knockout mice may contribute to augmented ingestive behaviors. Considering that pancreatic endocrine function is strongly influenced by circadian rhythms^44^, with hormone secretion and glucose handling varying across the light-dark cycle, the temporal specificity of this phenotype may reflect enhanced metabolic responsiveness at the onset of the active phase in the absence of MOPRs. In this context, improved glycemic regulation during this circadian window may facilitate the initiation of feeding bouts without altering meal size.

While our findings suggest that altered pancreatic function may contribute to the ingestive phenotype, an alternative or complementary explanation is that MOPRs in the brain may be driving these changes. As mentioned in the introduction, central MOPRs are well established regulators of feeding and reward related behaviors. MOPRs within mesolimbic and corticolimbic circuits including the ventral tegmental area (VTA) ^32–3^, nucleus accumbens (NAc)^45–48^, dorsal striatum^49^, amygdala^50–52^, and parabrachial nucleus (PBN)^32,53,54^ modulate the hedonic and motivational aspects of food intake. Thus, the increased meal frequency observed in global knockout males may reflect disruption of these brain circuits, rather than or in addition to, pancreatic mechanisms.

Unlike male knockout mice, female knockout mice did not differ in any metabolic assay tested compared to wild type female mice. Similarly, overall ingestive patterns were unaffected by MOPR deletion. However, while overall ingestive phenotypes were not different, further analyses indicated that, like male knockout mice, female knockout mice also increased the number of meals just prior to and at the onset of the dark cycle. While these phenotypes may be more subtle in female mice, it is possible that under certain conditions (e.g., time of day, stress), females may display phenotypes that would otherwise be overridden by counterregulatory processes.

The clear divergence between male and female knockout mice aligns with prior evidence that opioid receptor signaling interacts with sex hormones to shape energy balance and reward processes. For example, estrogen and progesterone have been shown to modulate opioid receptor function^55,56^, receptor activity^57^ and expression^58^. Such modulation may underlie the attenuated or context-dependent phenotypes observed in female knockouts, as fluctuating hormonal states could buffer against or compensate for receptor loss. Though not assessed here, future studies recording estrous cycle or manipulating estradiol/progesterone signaling could clarify whether sex hormones impact MOPR deletion on metabolic and motivational control of feeding.

A major question asked in our study was whether mu opioid receptors on pancreatic alpha cells could modulate metabolic or ingestive phenotypes. Consistent with prior work, male conditional deletion of MOPRs from alpha cells did not differ from controls in insulin or glucose tolerance, nor in body weight. Here, we additionally show that other metabolic biomarkers are similarly unchanged. Unlike constitutive knockout mice, conditional knockout of MOPRs from alpha cells in male mice did not alter total consumption over the testing period. However, male conditional knockout mice initiated significantly more meals than controls, an effect most pronounced during the light phase, and these increases were accompanied by smaller average meal sizes. This pattern partly mirrors the phenotype observed in global knockouts (i.e., increased meal frequency), but with a compensatory shift toward smaller meals that preserved total intake and body weight. Female conditional knockout mice showed a similar but more restricted phenotype. Like males, they consumed smaller meals across the circadian cycle, but this effect was not accompanied by differences in total intake, meal number, or body weight. Thus, both sexes demonstrated reduced meal size following α-cell MOPR deletion, while only males exhibited an increase in meal frequency.

One interpretation of our data is that loss of α-cell MOPRs produces an α-cell-centric dysregulation that alters ingestive microstructure through changes in glucoregulatory hormones rather than central appetitive drive. α-cells are the primary source of glucagon, a hormone that plays a key role in maintaining glucose homeostasis but also influences feeding behavior and post-prandial energy balance^59–61^. In addition to its classical role in stimulating hepatic glucose production, glucagon has been shown to suppress food intake and modulate satiety signaling, in part through hepatic–vagal pathways^62–66^ and effects on gastric emptying^67–69^. Through these mechanisms, glucagon signaling can influence the termination of meals and the interval between feeding bouts. Moreover, emerging evidence suggests that intra-islet signaling can influence systemic metabolic and behavioral responses, with α-cell derived peptides including glucagon and locally processed GLP-1 affecting satiety, gastric motility, and feeding microstructure. Consistent with this framework, peripheral modulation of MOPRs within pancreatic islets alters islet hormone secretion^39^, and disruption of MOPR signaling in α-cells could therefore shift glucagon dynamics. Such changes in endocrine signaling may alter satiety feedback following meals, providing a potential explanation for the altered meal patterning observed in conditional knockout mice, where decreased meal size is offset by increased meal frequency despite minimal changes in overall intake.

Our findings have broader implications for both metabolic disease and opioid research. In the context of diabetes and obesity, our data suggests that MOPR signaling influences not only glucose homeostasis but also the microstructural organization of feeding. Increased meal frequency and reduced meal size are patterns associated with altered glycemic control and may impact long-term risk for metabolic dysfunction^70,71^. Although α-cell MOPR deletion did not overtly affect glucose tolerance under baseline conditions, prior work has shown that conditional knockout mice are more susceptible to impaired glucose tolerance following a mild high-fat diet^39^, suggesting that MOPR loss may reveal metabolic vulnerabilities under physiological stress. The observed changes in feeding microstructure may represent early or compensatory adaptations with potential relevance to metabolic disease progression.

Opioids remain a cornerstone of treatment for painful diabetic neuropathy, yet the mechanisms by which chronic opioid exposure influences metabolic regulation are poorly understood. Our results suggest that both central and peripheral MOPR populations participate in the regulation of feeding and glycemia, raising the possibility that opioid-based treatments could inadvertently exacerbate or mitigate metabolic dysregulation in vulnerable populations. The dissociation between brain and pancreatic MOPRs highlights the importance of considering tissue-specific opioid actions, both in designing novel therapies and in predicting side effects of long-term opioid use.

## Acknowledgements

We thank David Piston for providing *GCG*-Cre x *Oprm1*^*fl/fl*^ mice. We thank the members of the Piston and Hughes Labs for their valuable feedback and support throughout this project. We are also grateful to the for their technical assistance.

## CRediT authorship contribution statement

**Daniel C. Castro**: Writing – review & editing, Supervision, Project administration, Methodology, Investigation, Funding acquisition, Conceptualization.

**Diego De Gregorio**: Writing – original draft, Visualization, Validation, Software, Resources, Formal analysis, Data curation.

---

**Fig. 1.**
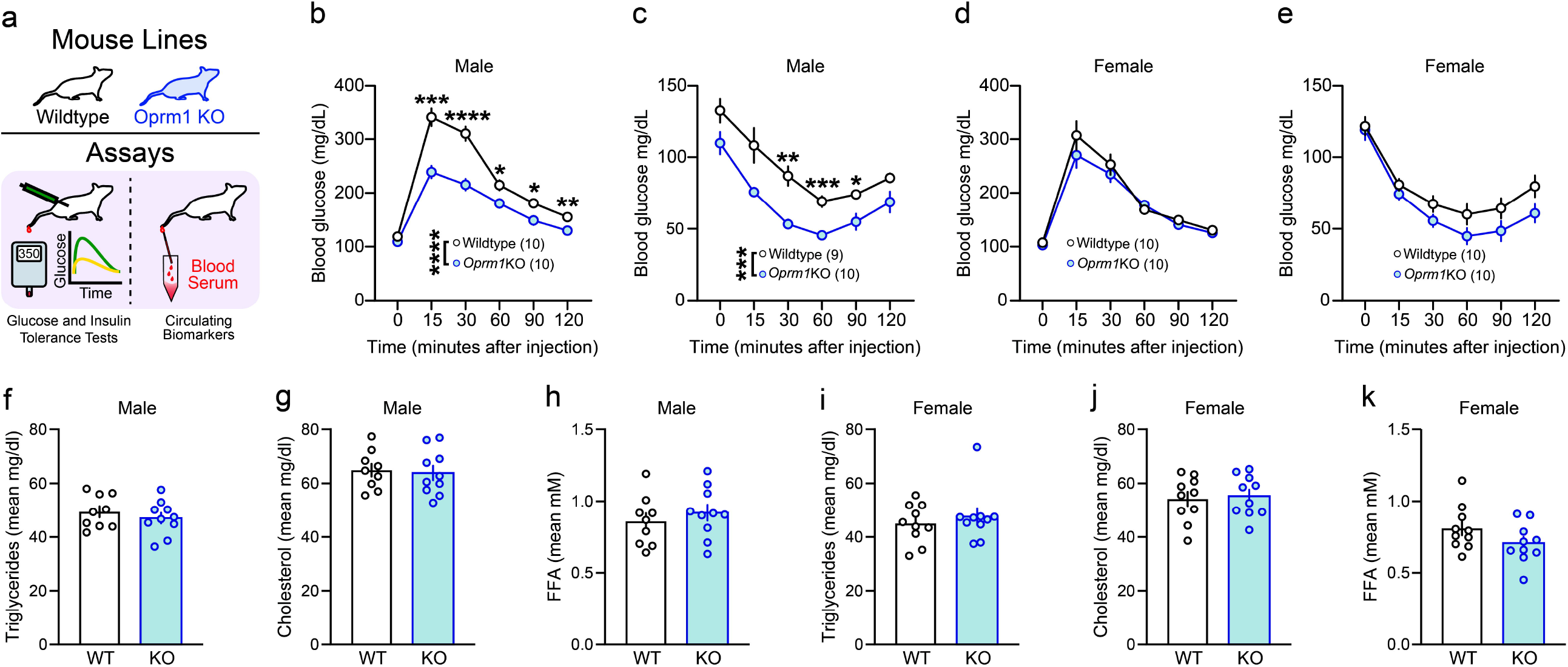
Male *Oprm1* KO mice show enhanced glucose tolerance and insulin sensitivity. (a) Schematic illustrating the mouse lines and metabolic assays used. (b, c) Intraperitoneal glucose tolerance test (GTT) and insulin tolerance test (ITT) in male wild type and *Oprm1* knockout mice (*Oprm1* KO or KO) mice. Male KO mice displayed enhanced glucose tolerance and increased insulin sensitivity relative to wild type controls. (d, e) GTT and ITT in female wild type and KO mice. No differences in glucose tolerance or insulin sensitivity were observed between female genotypes. (f-h) Serum triglycerides, cholesterol, and free fatty acids (FFA) in male mice. (i-k) Serum triglycerides, cholesterol, and FFA in female mice. No differences in serum biomarkers were detected in either sex. Data are presented as mean ± SEM. KO, knockout; GTT, glucose tolerance test; ITT, insulin tolerance test; FFA, free fatty acids.

**Fig. 2.**
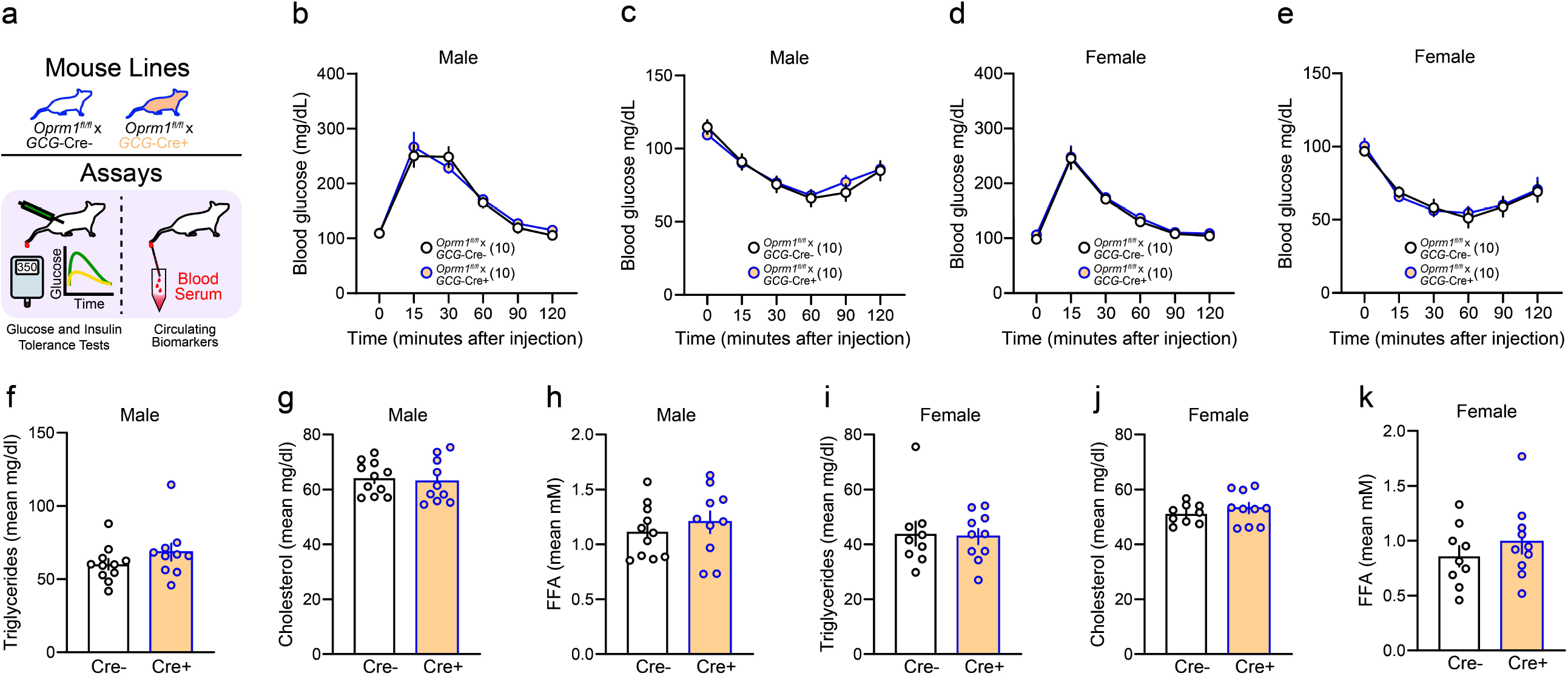
Male *Oprm1* KO mice show greater body mass and increased food consumption. (a) Schematic illustrating the mouse lines and ad libitum feeding assay. (b, c) Average body weight of male and female mice, respectively. (d) Average pellets consumed over the 72-hour testing period in male mice. (e) Average pellet consumption plotted in 4-hour intervals across the light/dark cycle in male mice. (f) Average number of meals and (g) their distribution across the light/dark cycle in male mice. (h) Average pellets per meal and (i) their distribution across the light/dark cycle in male mice. Male KO mice weighed more than wild type controls and consumed more pellets overall, with elevated consumption concentrated in the dark phase. (j) Average pellets consumed over the 72-hour testing period in female mice. (k) Average pellet consumption plotted in 4-hour intervals across the light/dark cycle in female mice. (i) Average number of meals and (m) their distribution across the light/dark cycle in female mice. (n)

**Fig. 3.**
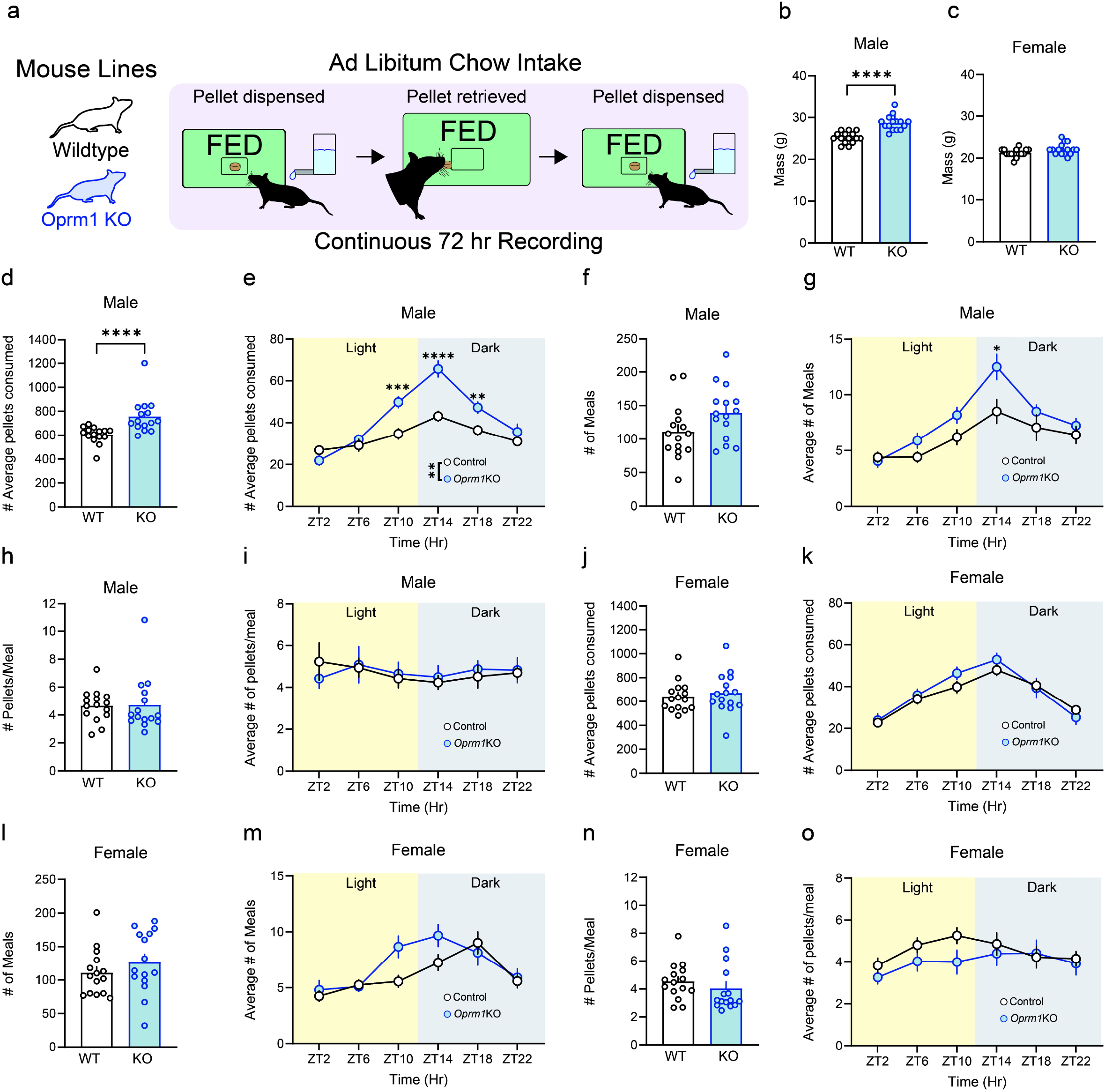
Metabolic phenotyping of *Oprm1*^*fl/fl*^ *x* GCG-Cre+ mice reveals no differences from Cre-controls. (a) Schematic illustrating the mouse lines and metabolic assays used. (b, c) Intraperitoneal glucose tolerance test (GTT) and insulin tolerance test (ITT) in male *Oprm1*^*fl/fl*^ *x GCG*-Cre mice (Cre+, Cre-). Male Cre+ mice did not differ from Cre-controls in glucose tolerance or insulin sensitivity. (d, e) GTT and ITT in female Cre+ or Cre-mice. Female Cre+ mice similarly showed no differences in glucose tolerance or insulin sensitivity relative to Cre-. (f–h) Serum triglycerides, cholesterol, and free fatty acids (FFA) in male mice. (i–k) Serum triglycerides, cholesterol, and FFA in female mice. No differences in serum biomarkers were detected in either sex. Data are presented as mean ± SEM. Cre+/-, conditional knockout; GTT, glucose tolerance test; ITT, insulin tolerance test; FFA, free fatty acids.

**Fig. 4.**
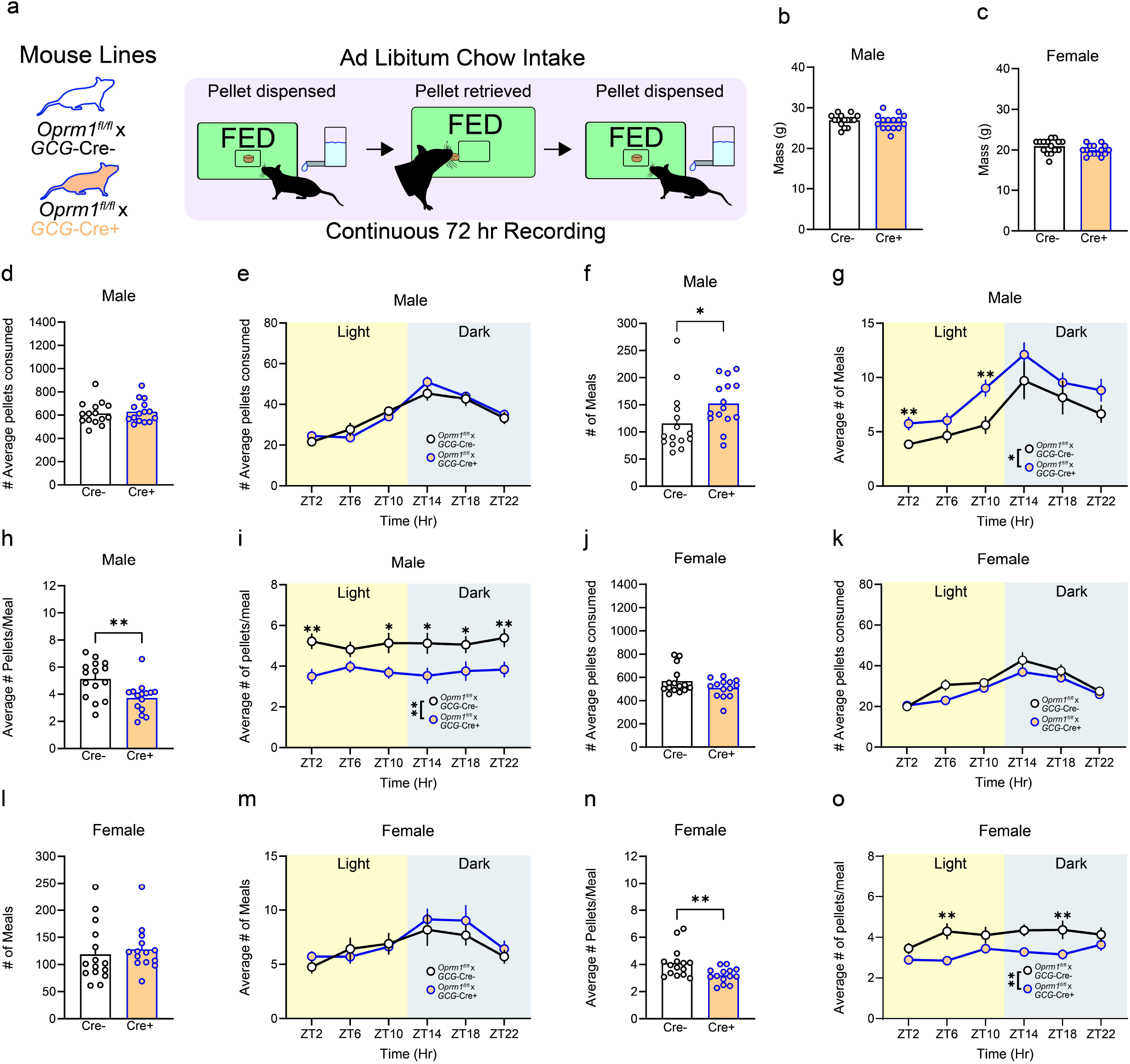
Alpha cell MOPR deletion alters ingestive microstructure without affecting total intake. (a) Schematic illustrating the mouse lines and ad libitum feeding assay used. (b, c) Average body weight of male and female mice, respectively. (d) Average pellets consumed over the 72-hour testing period in male mice. (e) Average pellet consumption plotted in 4-hour intervals across the light/dark cycle in male mice. (f) Average number of meals and (g) their distribution across the light/dark cycle in male mice. (h) Average pellets per meal and (i) their distribution across the light/dark cycle in male mice. Male Cre+ mice did not differ from Cre-controls in body weight or total intake, though Cre+ males ate more meals of smaller size, an effect most prominent during the light phase. (j) Average pellets consumed over the 72-hour testing period in female mice. (k) Average pellet consumption plotted in 4-hour intervals across the light/dark cycle in female mice. (l) Average number of meals and (m) their distribution across the light/dark cycle in female mice. (n) Average pellets per meal and (o) their distribution across the light/dark cycle in female mice. Female Cre+ mice did not differ from Cre-controls in body weight, total intake, or meal number, though Cre+ females consistently had smaller meal sizes regardless of the light/dark cycle. Data are presented as mean ± SEM. aKO, Cre+/-, conditional knockout; ZT, Zeitgeber time.

